# Decomposing spatial heterogeneity of cell trajectories with Paella

**DOI:** 10.1101/2022.09.05.506682

**Authors:** Wenpin Hou, Zhicheng Ji

## Abstract

Spatial transcriptomics provides a unique opportunity to study continuous biological processes in a spatial context. We developed Paella, a computational method to decompose a cell trajectory into multiple spatial sub-trajectories and identify genes with differential temporal patterns across spatial sub-trajectories. Applied to spatial transcriptomics datasets of cancer, Paella identified spatially varying genes associated with tumor progression, providing insights into the spatial heterogeneity of cancer development.

## Main

Cell trajectory inference methods^1–5^ enable the study of continuous biological processes by computationally ordering cells based on their gene expression profiles measured by single-cell RNA-sequencing (scRNA-seq)^6^. A major limitation of scRNA-seq is that cellular spatial information is lost during tissue dissection. Thus, it is unknown whether the inferred cell trajectory may arise from different spatial locations and whether there are gene expression variations across spatial sub-trajectories. Such knowledge is crucial in deciphering the spatial heterogeneity of cell development and differentiation processes.

Spatial transcriptomics profiling measures both the gene expression and the spatial locations of cells^7–9^, offering the potential to decompose the spatial heterogeneity of cell trajectories. However, most existing cell trajectory methods such as Monocle^1, 2^, Slingshot^4^, and TSCAN^3^ are designed for scRNA-seq data and do not consider spatial information. Methods for spatial trajectory analysis such as SpaceFlow^10^, SPATA^11^, and SpatialPCA^12^ infer and visualize pseudotime trajectories on a spatial map but do not study the spatial heterogeneity of a cell trajectory. stLearn^13^ is able to identify spatial sub-trajectories but has several limitations. First, it can only detect simple trajectory patterns transitioning from one cell cluster to another, but it cannot detect complicated patterns such as concentric rings. Second, a single spatial trajectory inferred by stLearn may appear in discrete spatial regions far away from each other, which is against the continuous nature of cell development and differentiation. Third, stLearn lack the functionality to systematically analyze differential dynamic gene expression patterns across spatial trajectories. Finally, stLearn cannot analyze manually defined cell trajectories. In summary, how to study the spatial heterogeneity of cell trajectories is a question yet to be answered.

To this end, we developed Paella to decompose the s<u>pa</u>tial h<u>e</u>terogeneity of ce<u>ll</u> tr<u>a</u>jectories (Figure 1a). Spatial decomposition helps explain the intrinsic variability of cell development processes and may ultimately improve the understanding of how heterogeneous functions and characteristics of different spatial regions are formed in the same tissue. Paella requires as input the spatial locations of cells or spatial spots and the cell trajectory information. Cell trajectories can be inferred by only using gene expression (e.g., Monocle^1, 2^, TSCAN^3^), by using both gene expression and spatial information (e.g., SpaceFlow^10^), or by some customized procedures (see the breast cancer example below). Paella then identifies a parsimonious set of spatially continuous sub-trajectories where each sub-trajectory represents a unidirectional process of cell progression. Finally, Paella identifies and analyzes genes with differential dynamic patterns across spatial sub-trajectories. These genes may serve as factors driving the spatial heterogeneity of cell trajectories.

**Figure 1.**
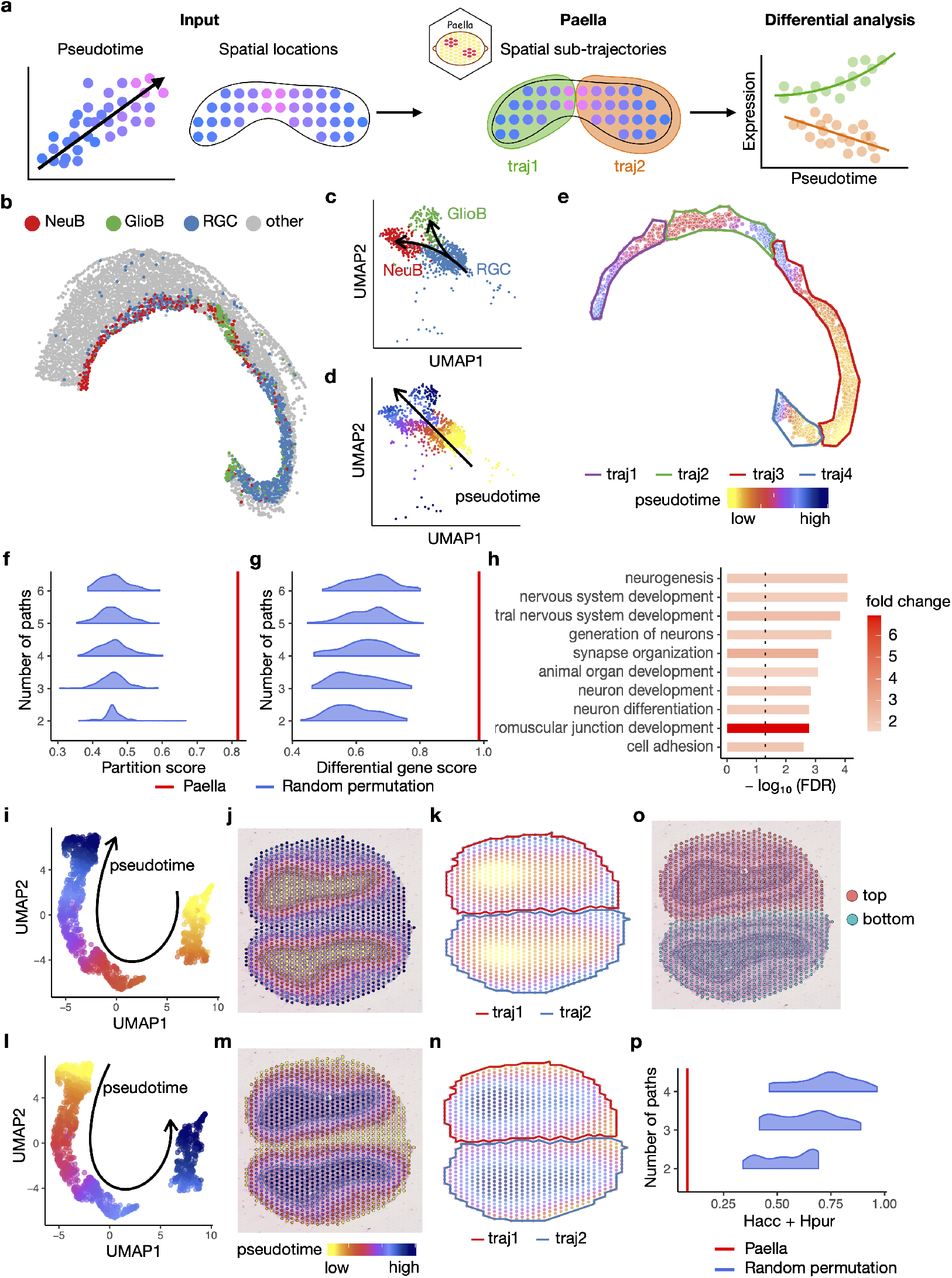
Paella schematic (**a**), analysis of mouse E14.5 midbrain data (**b**-**h**), and analysis of olfactory bulb data (**i**-**p**). **a**, A schematic view of Paella algorithm. **b**, Spatial locations of RGCs, NeuBs, GlioBs, and other cells. **c**, Cell trajectories of RGCs differentiating into NeuBs and GlioBs shown on Uniform Manifold Approximation and Projection^44^(UMAP) space. **d**, Pseudotime values and direction shown on UMAP space. **e**, Spatial sub-trajectories identified by Paella. A sub-trajectory is abbreviated as a “traj” on the plot. **f**, Partition scores comparing Paella with random permutation. **g**, Differential gene scores comparing Paella with random permutation. **h**, Top GO terms enriched in the spatially differential genes reported by Paella. An FDR cutoff of 0.05 is indicated as a vertical black dashed line. **i**,**l**, Pseudotime values and direction shown on UMAP space. **j**,**m**, Pseudotime values shown on H&E plot. **k**,**n**, Spatial sub-trajectories identified by Paella. **i-k**, Pseudotime goes from inner to outer rings. **l-n**, Pseudotime goes from outer to inner rings. **o**, Manual annotation of two olfactory bulbs. **p**, Partition agreement comparing Paella and random permutation. The lower the better.

As a proof of concept, we applied Paella to a mouse E14.5 dorsal midbrain spatial transcriptomics dataset generated by Stereo-seq^14^ (Figure 1b). The dataset contains radial glia cells (RGCs), neuroblasts (NeuBs), and glioblasts (GlioBs). RGCs differentiate into two branches of NeuBs and GlioBs (Figure 1c). RGCs are enriched in two spatial regions, while NeuBs and GlioBs occupy spatial regions neighboring RGCs (Figure 1b), representing tubular patterns of spatial transitions between cell types. For evaluation purposes, we merged the two branches into one (Figure 1d) and evaluated whether Paella is able to identify spatial sub-trajectories that reflect the differences between NeuBs and GlioBs. Paella identifies four spatial sub-trajectories, each originating from a region enriched in RGCs to a region enriched in NeuBs or GlioBs (Figure 1e), which agrees with the spatial locations of the three cell types. Due to the tubular nature of this dataset, we evaluated if each spatial sub-trajectory represents a unidirectional transition using a partition score (Methods). The partition score measures if a pair of cells with similar pseudotime values are spatially close to each other. Paella reaches a partition score of 0.817 and performs better than a benchmarking method that randomly permutes the sub-trajectory labels (Figure 1f, Methods). To evaluate if Paella recovers the gene expression differences between NeuBs and GlioBs, we compared the overlap between genes with differential temporal patterns across Paella spatial sub-trajectories and genes with differential temporal patterns between actual NeuB and GlioB trajectories (Methods). Paella reaches a score of 0.871 and again performs better than the benchmarking method (Figure 1g). The differential genes identified by Paella are enriched in gene ontology (GO) terms such as neurogenesis and nervous system development process, consistent with the biological process of neuron development (Figure 1h).

Next, we demonstrate Paella’s ability to analyze more complex spatial structures using a spatial transcriptomics dataset of a mouse olfactory bulb (OB) generated by 10x Visium Spatial Transcriptomics technology^15^. The anatomical structure of OB consists of multiple layers of tissues representing concentric rings^16^. Using gene expression, we constructed a trajectory from inner rings to outer rings (Figure 1i,j), and Paella identifies two spatial sub-trajectories corresponding to the two OBs in the tissue slice (Figure 1k). We then reversed the trajectory direction and Paella is still able to detect the same two spatial sub-trajectories (Figure 1l-n). To evaluate the performance, we manually annotated the spatial spots belonging to the two OBs (Figure 1o). Paella’s spatial sub-trajectory assignments agree well with the manual annotations and Paella outperforms the benchmarking method (Methods, Figure 1p).

Then, we applied Paella to study the spatial heterogeneity of tumor progression. We first analyzed a human glioblastoma (GBM) spatial dataset generated by Visium^17^. We inferred a pseudotime trajectory of glioblastoma stem cell (GSC) differentiation. Along this trajectory, GSC marker genes *SOX2*^18^ and *OLIG2*^19^ show decreasing expression and tumor proliferation and progression markers *VEGFA*^20^ and *CXCL8*^21^ show increasing expression (Figure 2a-b, Supplementary Figure 1). We applied Paella and identified two spatial sub-trajectories with different origins (Figure 2c). Paella identified 5321 out of 12905 genes (41.2%) with differential temporal patterns across the two spatial sub-trajectories (Figure 2d-e), indicating that spatial heterogeneity is associated with a large variation in transcriptomic programs. In addition to identifying 4251 genes with mean changes, Paella also identified 3128 genes with trend changes which can only be detected by pseudotime analysis. Many of the top differential genes have been reported to be linked to glioblastomia, including *PRRX1*^22^, *MRC2*^23^, *LAMC1*^24^, *BNIP3*^25^, *VEGFA*^26^, *SLC2A1*^27^, and *NTRK2*^28^. Many enriched GO terms associated with the differential genes (Figure 2f) are reported in previous literature, including blood vessel development^29^, chemical synaptic transmission^30^, and ion channels^31^. Utilizing the gene expression data from The Cancer Genome Atlas (TCGA^32^, Methods), we found that the spatially differential genes identified by Paella have the highest overlap with highly variable genes across GBM and low-grade glioma (LGG) samples from TCGA, followed by samples of other cancers types (Figure 2g). This indicates that Paella is able to identify spatially differential genes relevant to the biological context, and the spatial variation provides a likely explanation of the variation across cancer samples. The overlap of Paella is also higher than the benchmarking method (Figure 2h).

**Figure 2.**
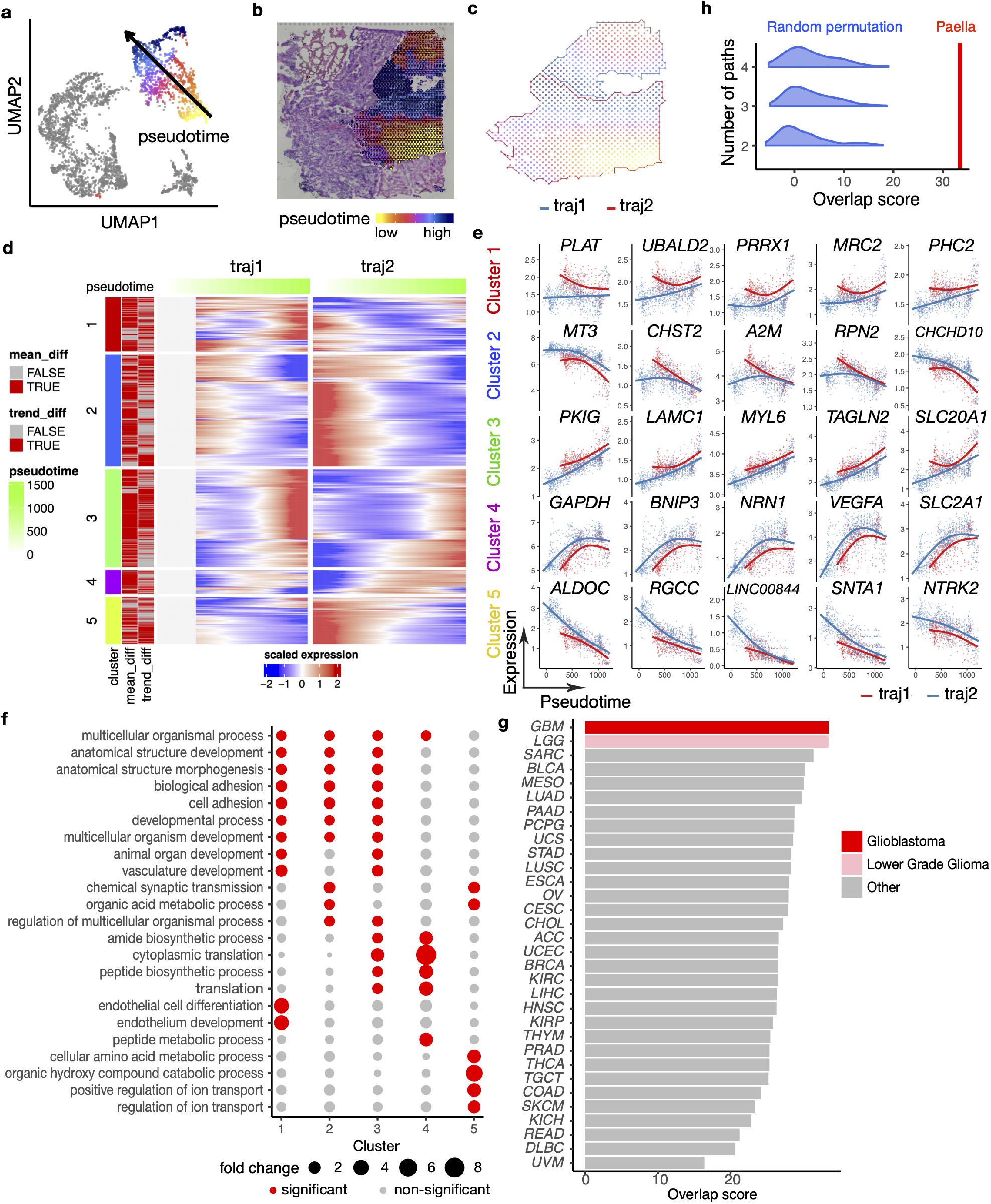
Paella analysis of human glioblastoma data. **a**, Pseudotime values and direction shown on UMAP space. **b**, Pseudotime values shown on H&E image. **c**, Spatial sub-trajectories identified by Paella. **d**, A heatmap showing all genes with differential temporal patterns across spatial sub-trajectories. The three columns on the left show the cluster of a gene and if the gene is mean or trend differential. **e**, Gene expression values and fitted GAM^45^ curves for the top five differential genes in each gene cluster. **f**, Top GO terms enriched in the differential genes. **g**, Overlap score of spatially differential genes identified by Paella and highly variable genes identified in TCGA^32^ samples with different cancer types. **h**, Overlap score comparing spatially differential genes identified by Paella or random permutation and highly variable genes identified in TCGA^32^ glioblastoma samples.

Finally, we applied Paella to a human breast cancer spatial dataset generated by Visium^33^. A unique feature of Paella is its flexibility of analyzing cell trajectories obtained by customized procedures, which is useful for analyzing spatial data manually annotated by biologists and pathologists. As an example, instead of inferring pseudotime from gene expression, we manually identified six carcinoma *in situ* regions and their centers according to a previous study^34^, and defined the pseudotime value of a spot as its distance to the nearest center of carcinoma *in situ* region on the spatial map (Figure 3a). The pseudotime represents cancer cells progressing from the center of the tumor to its border. We then applied Paella and identified six spatial sub-trajectories (Figure 3b), agreeing with the six carcinoma *in situ* regions. Paella identified 3105 out of 13643 genes (22.8%) with differential temporal patterns across spatial sub-trajectories (Figure 3c-d), again showing a large number of genes associated with spatial heterogeneity of tumor progression. Many of the top differential genes have been reported to be linked to breast cancer, including *CALML5*^35^, *ABCC5*^36^, *KLF6*^37^, *NR4A1*^38^, *SPDEF*^39^, and *MRPS30-DT*^40^. We again found many enriched GO terms (Figure 3e) to be reported in previous literature, such as immune and antigen presenting^41^, blood vessel development^42^, and cell adhesion^43^. Similar to GBM, the spatially differential genes identified by Paella have the highest overlap with highly variable genes across *BRCA* (breast cancer) samples in TCGA^32^ (Figure 3f). Within breast cancer subtypes, the overlap is highest with Luminal A, which agrees with the classification of this spatial sample (Supplementary Figure 2). The overlap of Paella is also higher than the benchmarking method for all TCGA^32^ breast cancer samples (Figure 3g) and Luminal A samples (Figure 3h).

**Figure 3.**
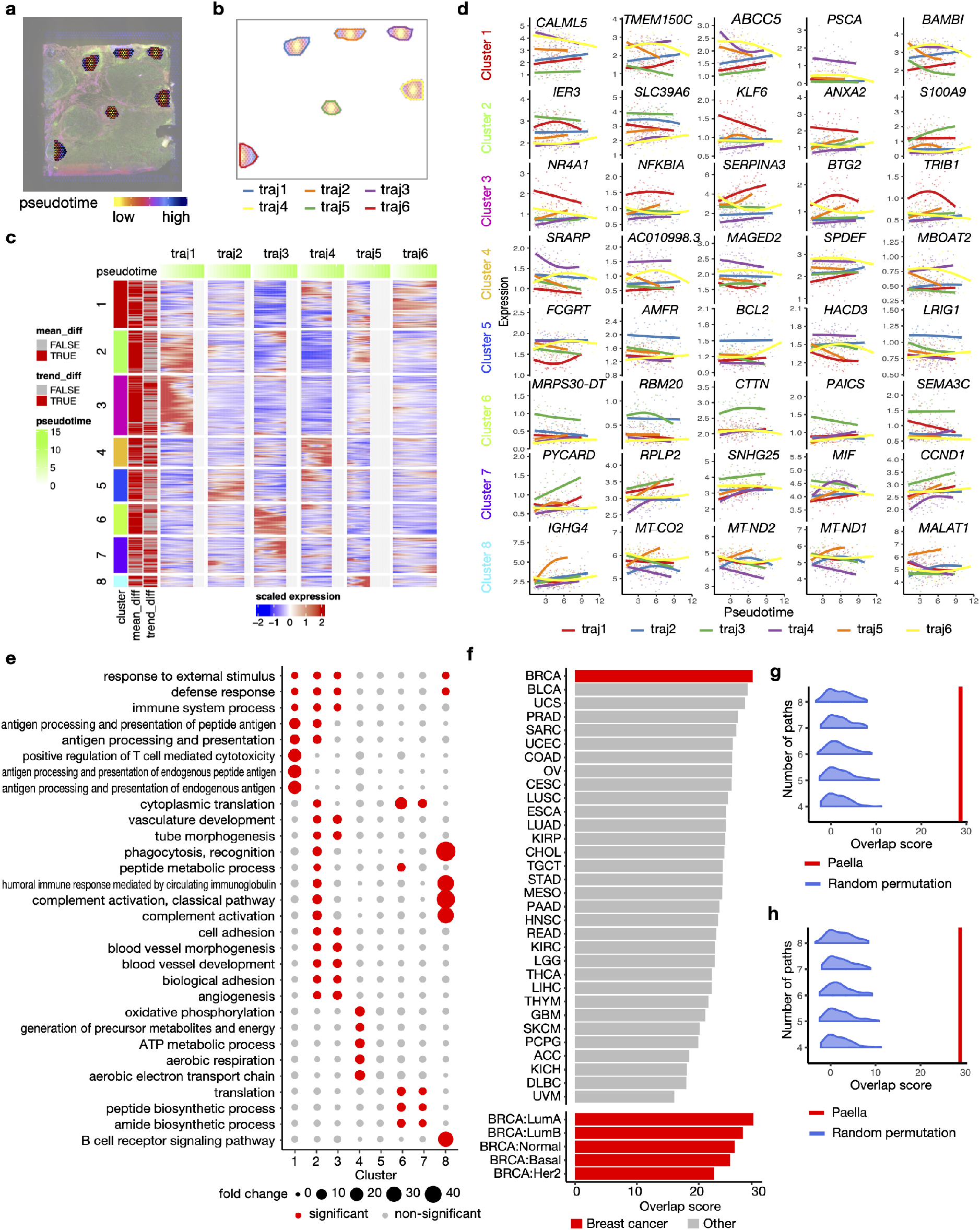
Paella analysis of human breast cancer data. **a**, Pseudotime values shown on immunofluorescent image. **b**, Spatial sub-trajectories identified by Paella. **c**, A heatmap showing all genes with differential temporal patterns across spatial sub-trajectories. The three columns on the left show the gene clusters and the gene’s differential pattern (mean or trend). **d**, Gene expression values and fitted gam curves for top 5 differential genes of each gene cluster. **e**, Top GO terms enriched in the differential genes. **f**, Overlap score comparing spatially differential genes identified by Paella and highly variable genes identified in TCGA^32^ samples with different cancer types (top) or with different breast cancer subtypes (bottom). **g-h**, Overlap score comparing spatially differential genes identified by Paella or random permutation method and highly variable genes identified in all TCGA^32^ breast cancer samples (**g**) or breast cancer samples with Luminal A subtype (**h**).

The genes with differential temporal patterns across spatial sub-trajectories identified by Paella from GBM and breast cancer datasets share a statistically significant overlap (*p*-value= 6.54 *×*10^*−* 87^, Supplementary Figure 3, Methods). The enriched GO terms of the overlapping genes include blood vessel development, cell adhesion, and responses to chemicals and stimulus (Supplementary Figure 4). This indicates that the spatial heterogeneity of tumor progression in different cancer types share some common gene pathways, which is worth further investigation with future spatial transcriptomics data in other cancer types.

## Methods

### Paella algorithm to identify spatial sub-trajectories

#### Overview

Paella is designed for spatial transcriptomics data with gene expression profiles and spatial locations of the cells, including data generated by spot-based technologies (e.g., Visium^15^) or FISH (fluorescence *in situ* hybridization) technologies (e.g., MERFISH^8^). Hereinafter, we use a node to refer to the smallest unit in the data, which is a spatial spot in spot-based technologies or a cell in FISH technologies.

The goal of Paella is to identify a parsimonious set of spatial sub-trajectories that are spatially continuous and represent unidirectional progressions of cell trajectory. For a spatial region with a tubular pattern, this means the cell trajectory progresses from a single start point to a single end point. For a spatial region with a concentric ring pattern, this means the cell trajectory progresses from outer rings to inner rings or inner rings to outer rings, depending on the direction of the cell trajectory.

Paella starts by constructing an undirected Delaunay network of nodes using their spatial locations. It then filters out isolated nodes far away from most other nodes, and filters out long edges. For each connected component of the Delaunay network, Paella converts the undirected network into two directed networks by comparing the pseudotime values of the two nodes connected by an edge, and identifies with three modes all node sets where nodes in each set are reachable from a common ancestry. Paella then sequentially assembles node sets that lead to the largest size of the union of all selected sets in each iteration. Each selected node set is a tentative spatial sub-trajectory. Nodes assigned to no or more than one sub-trajectories are reassigned to create final spatial sub-trajectories. Finally, the mode that produces a parsimonious set of unidirectional sub-trajectories is selected to generate the final spatial sub-trajectories.

Below are details of the Paella algorithm in sequential order.

#### Input

Paella requires two inputs: (1) the pseudotime values for all *S* nodes, denoted as *t*_*i*_ ∈ R for node *i*, and (2) the *d*-dimensional spatial coordinates for all *S* nodes, denoted as 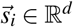 for node *i*. All datasets analyzed in this study have 2-dimensional spatial coordinates (*d* = 2), but Paella can also deal with datasets with 3-dimensional spatial coordinates (*d* = 3).

#### Delaunay triangulation network

Similar to `Giotto`^46^, Paella builds a Delaunay triangulation network (`geometry::delaunay()`) that identifies edges for neighbouring nodes. The Delaunay network serves as the basis of multiple subsequent tasks of Paella. We denote the constructed network as *G*(*V, E*), where *V* is the set of all nodes and *E* is the set of all edges in the graph.

#### Removal of isolated nodes

Paella removes isolated nodes that are far away from all other nodes. We define the weight of an edge in *G* as 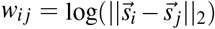, which is the log of Euclidean distance between two nodes *i* and *j* connected by this edge. For a pair of nodes *k* and *l*, Paella then calculates 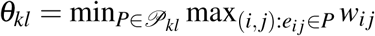, where *𝒫*_*kl*_ is the set of all paths connecting *k* and *l*, and each path is a set of edges. *e*_*i j*_ is an edge connecting nodes *i* and *j* in the network. This is similar to the bottleneck problem but in a reversed direction. We further set θ_*kk*_ = 0. For any node *k*, Paella then calculates an isolation score *η*_*k*_ = ∑_*l*_ θ_*kl*_ /*S*. The isolation score represents the overall level of isolation of this node from all other nodes.

To filter out nodes with very large isolation scores, Paella fits a density curve of all isolation scores (`stats::density()`). Paella identifies a set of values *A*, which includes min *η*_*k*_ *−* 1, max *η*_*k*_ + 1, and all valley points with negative left derivatives and positive right derivatives on the density curve. All elements in *A* are sorted in an increasing order. The *l*-th element in the sorted set *A* is denoted as *A*_*l*_. The region between two consecutive valley points are considered as a peak. The number of nodes falling in peak between *A*_*l*_ and *A*_*l*+1_ is counted as ∑_*k ∈ S*_ *I*(*A*_*l*_ < *η*_*k*_ *≤ A*_*l*+1_). Suppose the peak between *A*_*l*_ and *A*_*l*+1_ is the first peak having less than 1% of total nodes, all nodes with *η*_*k*_ > *A*_*l*_ are filtered out. If no peak has less than 1% of total nodes, no node is excluded.

#### Removal of long edges

Similar to the previous step, Paella uses a density peak approach to filter out edges with large *w*_*i j*_.

#### Identifying network connected components

After filtering out nodes and edges, the original network may not remain fully connected. Paella identifies all connected components in the network, and all subsequent operations are performed within each connected component separately. The final set of spatial sub-trajectories is the union of spatial sub-trajectories identified in all connected component.

#### Smoothing pseudotime value

Paella first smooths nodes whose pseudotime values are very different from the pseudotime values of neighbouring nodes. For a node *i*, Paella calculates a pseudotime neighbouring difference 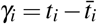 Here 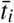 is the averaged pseudotime values of all nodes sharing edges with *i* in the network. For a node *i* with |*γ*_*i*_| larger than 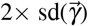 (sd is the standard deviation, 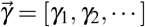, its original pseudotime value *t*_*i*_ is replaced with 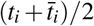. The pseudotime values for all other nodes remain unchanged.

Then Paella fits a generalized addictive model (GAM)^45^ of pseudotime values against the spatial coordinates (`mgcv::gam()`), and uses the fitted value 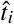 to replace the original pseudotime values *t*_*i*_.

#### Converting undirected network to directed network

To prepare for following steps, Paella converts the undirected Delaunay network into two directed networks. Paella first calculates a pseudotime tolerance value. The original pseudotime values before smoothing are randomly permuted and the GAM model is refitted with the permuted values. For each pair of nodes connected by a network edge, a difference of their fitted pseudotime values is calculated. A standard deviation is calculated of all such differences across all connected node pairs. The whole permutation process is repeated for 100 times, and the pseudotime tolerance value *T* is set as 2 *× the median of these* 100 *standard deviations*.

To create the first directed network *G*_1_, each undirected edge connecting nodes *i* and *j* in the original network are replaced with two directed edges: *i → j* and *j → i*. An edge *i → j* is retained if 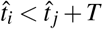. The second directed network *G*_2_ is created using the same procedure but with the retaining criteria of 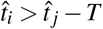.

#### Identifying progression sets

Paella identifies progression sets on the two directed networks *G*_1_ and *G*_2_. A progression set is a set of nodes that are spatially connected and represent unidirectional cell progression. The progression sets are identified in three modes. In the first mode, a progression set primed by node *i* is defined as the set of nodes reachable from node *i* in network *G*_1_. A set of all progression sets *O*_1_ is identified by going through all nodes as priming nodes in the network. In the second mode, a set of all progression sets *O*_2_ is identified on network *G*_2_ using the same procedure as the first mode. In the third mode, the progression sets are identified as *O*_3_ = {*M ∩ N M* | *∈ O*_1_, *N ∈ O*_2_}

Paella then filters out progression sets with very small numbers of nodes. Based on a binomial distribution, a progression set size cutoff is defined as 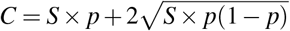. For the first two modes, *p* is the proportion of all possible pairs of nodes that are connected on *G*_1_ (denoted as *p*_1_) or *G*_2_ (denoted as *p*_2_). For the third mode, *p* = *p*_1_ *× p*_2_. All progression sets with numbers of nodes smaller than *C* are filtered out.

#### Creating spatial sub-trajectories

Paella sequentially assembles progression sets to create initial sub-trajectories. For the set of progression sets obtained in the first mode *O*_1_, Paella iteratively selects the progression set that leads to the largest size of the union of all progression sets selected at that iteration. Specifically, for iteration *i*, Paella selects a progression set *K*^(*i*)^ from *O*_1_ that maximizes |*K*^(1)^ ∪ *…* ∪ *K*^(*i*)^ |. Here | | is the cardinality of a set. The iteration stops at *i* when |*K*^(1)^ ∪ *…* ∪ *K*^(*i*+1)^ |*−* | *K*^(1)^ ∪ *…* ∪ *K*^(*i*)^ | < *C*. Each progression set now becomes an initial spatial sub-trajectory. To reconcile undetermined nodes belonging to zero or more than one initial sub-trajectories, each of such undetermined node *i* is reassigned the sub-trajectory identity of the determined node (node with exactly one initial sub-trajectory assignment) closest to *i* on the network.

The procedure is repeated on progression sets obtained from each of the three modes.

#### Determining mode

The three modes generate three sets of spatial sub-trajectories that can be potentially different. Paella evaluates if a mode creates parsimonious and unidirectional spatial sub-trajectories, and selects the optimal mode to create final spatial sub-trajectories. A set of spatial sub-trajectories is parsimonious if no pair of spatial sub-trajectories can be merged into a single unidirectional sub-trajectory. A sub-trajectory is unidirectional if for any pair of nodes there exists a path linking the two nodes and the range of the pseudotime of all nodes along this path does not substantially exceed the range of pseudotime of the two end points.

Specifically, Paella defines a traverse cost on the undirected Delauney network for each pair of nodes *i* and *j* as 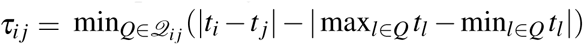. Here *𝒬*_*i j*_ is the set of all paths connecting *i* and *j*, and each element in *𝒬*_*i j*_ is a set of nodes on one path. However, calculating *τ*_*i j*_ for all pairs of nodes in the network is computationally intractable. So Paella applies an approximate algorithm to obtain 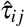 which is close to *τ*_*i j*_. The algorithm is similar to the Dijkstra’s algorithm^47^. For a root node *i*, the weight of any node *j* is set to be | *t*_*i*_*− t* _*j*_|. The algorithm initializes the set of visited nodes as the root node, and the set of unvisited nodes as all other nodes. In each iteration, the algorithm visits the unvisited node that is connected with visited nodes and has the smallest node weight, and records the visiting path. The iteration stops when all nodes are visited. The algorithm constructs a new graph *G*^*′*^ where all nodes are the vertices and the visiting paths are the edges. Finally, the algorithm calculates 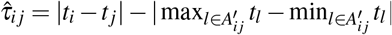 where 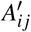 is the path connecting *i* and *j* on *G*^*′*^.

Paella then identifies pairs of neighbouring spatial sub-trajectories *ℒ*_1_ and *ℒ*_2_. *ℒ*_1_ and *ℒ*_2_ are spatially neighbouring if there exists at least one edge linking one node in *ℒ*_1_ and one node in *ℒ*_2_. For each pair of neighbouring sub-trajectories *ℒ*_1_ and *ℒ*_2_, a traverse cutoff is determined as mean + 2 *×* sd of the traverse scores 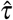 for all pairs of nodes that are either both within *ℒ*_1_ or both within *ℒ*_2_. A traverse proportion is defined for the pair *ℒ*_1_ and *ℒ*_2_ as the proportion of all cross-trajectory node pairs (one node is in *ℒ*_1_ and the other is in *ℒ*_2_) with traverse scores larger than the traverse cutoff. The traverse proportion is calculated for each pair of neighbouring sub-trajectories.

Finally, three metrics are calculated for each mode: (1) *I*(*ω*), *ω* = {the smallest traverse proportion is larger than 0.05}, (2) the total number of neighbouring sub-trajectory pairs, and (3) the averaged traverse proportion across all neighbouring sub-trajectory pairs. The three modes are ordered sequentially and decreasingly by the three metrics. If the first scores are tied, modes are ordered by the second scores. If the second scores are tied, modes are ordered by the third scores. The mode with the top order (largest score) is selected as the best mode.

### Differential gene analysis across spatial sub-trajectories

#### Identifying differential genes across spatial sub-trajectories

Similar to a previous study^48^, Paella identifies three types of differential genes: overall differential, mean differential, and trend differential. For each gene, three regression models are fit using the generalized additive model (GAM) in R’s `mgcv` package^45^. The formulas for the three models are listed below:

M1: *g* ~ *s*(*t*,*k* = 3)
M2: *g* ~ *α* +*s*(*t*,*k* = 3)
M3: *g* ~ *α* +*s*(*t*,*by* = *α*,*k* = 3)

Here *g* is a gene expression vector for a gene, *α* is a categorical vector of spatial sub-trajectory assignments, and *t* is a vector of pseudotime values. M1 is a baseline model that fits a universal curve for all sub-trajectories. M2 is a model allowing a mean shift that fits multiple parallel curves for different sub-trajectories. M3 is a model allowing both mean and trend shift fitting multiple curves that may not be in parallel with each other for different sub-trajectories.

Multiple likelihood ratio tests are performed between pairs of the three models. M1 and M3 are compared to test for overall differential. M1 and M2 are compared to test for mean differential. M2 and M3 are compared to test for trend differential. For each type of differential test, *p*-values are adjusted for multiple testing using the BH procedure^49^.

#### Gene clustering

For each gene, a GAM model is fitted separately for each spatial sub-trajectory with the formula *g∼ s*(*t, k* = 3) in `mgcv`, where *g* is the gene expression and *t* is the pseudotime for nodes within that spatial sub-trajectory. For each sub-trajectory, fitted values are obtained on 200 equidistant pseudotime values across the full pseudotime range. Values that are outside the range of the pseudotime of this sub-trajectory are excluded. For each gene, the fitted values across all sub-trajectories are pooled and standardized to have mean 0 and standard deviation of 1. Hierarchical clustering (complete agglomeration) is then performed on the pooled and standardized fitted gene expression values. By default, `findPC`^50^ is used to automatically select the number of gene clusters based on the proportion of within cluster sum of squares and total sum of squares (maximum of 20 clusters).

#### GO enrichment analysis

GO enrichment is performed with the `topGO` R package^51^, and *p*-values are adjusted for multiple testing using the BH procedure^49^.

### Spatial data processing and cell trajectory inference

#### Spatial data of mouse E14.5 dorsal midbrain

The processed gene expression matrix, spatial locations, and pseudotime values of mouse E14.5 dorsal midbrain cells were directly downloaded from the orginal paper (https://db.cngb.org/stomics/mosta/)^14^. Only NeuBs, GlioBs, and RGCs were retained. The processed gene expression data were imputed by `SAVER`^52^ for differential gene analysis.

#### Spatial data of mouse olfactory bulb

The original gene expression count matrix, spatial locations of spots, and the H&E image of a 10x Visium mouse olfactory bulb dataset were downloaded from 10x website^15^. Spots with more than 500 number of reads were retained. The gene expression data were processed using SCTransform^53^. Dimension reduction and clustering were then performed using standard Seurat pipeline^54^. Cell trajectory was obtained by TSCAN^3^ on UMAP space using default settings.

#### Spatial data of human glioblastoma

The original gene expression count matrix, spatial locations of spots, and the H&E image of a 10x Visium human glioblastoma dataset were downloaded from the 10x website^17^. Spots with more than 500 reads were retained. The gene expression data were processed using `SCTransform`^53^. Dimension reduction and clustering were performed using default `Seurat` pipeline^54^. Cell trajectory was inferred on a subset of cells with high expression of OLIG2 and SOX2 or VEGFA and CXCL8, and the pseudotime values were obtained by `TSCAN`^3^ on UMAP space with default settings. The original count data were imputed by `SAVER`^52^ for differential gene analysis.

#### Spatial data of human breast cancer

The original gene expression count matrix, spatial locations of spots, and the immunofluorescent image of a 10x Visium human breast cancer dataset were downloaded from 10x website^33^. Spots with more than 500 number of reads were retained. The gene expression data were processed using SCTransform^53^. Dimension reduction and clustering were then performed using standard Seurat pipeline^54^. Pseudotime values were obtained using a manual procedure described above. The original count data were imputed by `SAVER`^52^ and the imputed values were used for differential gene analysis.

### TCGA data processing and analysis

Gene expression data of all cancer types were downloaded from TCGA^32^ using the R/Bioconductor^55^ package `TCGAbiolinks`^56^. Only samples from primary tumor were retained and FFPE samples were excluded. Unstranded TPM values were log_2_ transformed after adding a pseudo count of 1. For multiple samples belonging to the same patient, their gene expression values were averaged to create a single gene expression vector for that patient.

### Evaluation

#### Benchmarking method

To benchmark the performance of Paella, we created a random permutation of sub-trajectory assignments of nodes with different numbers of sub-trajectories. To generate a random permutation for *k* sub-trajectories, we first sampled a vector of probabilities *p*_*i*_ from a Dirichlet distribution with *k* categories 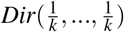. Then we randomly assigned nodes to sub-trajectories. The number of nodes in *i*-th random sub-trajectory is *Sp*_*i*_ where *S* is the total number of nodes. If *Sp*_*i*_ < 20 for *i*-th sub-trajectory, we reassign 20*− Sp*_*i*_ nodes from the sub-trajectory with the largest number of nodes to the *i*th sub-trajectory, so that each sub-trajectory has a minimum of 20 nodes.

#### Partition score in mouse E14.5 dorsal midbrain data

Suppose there are *k* spatial sub-trajectories. For each sub-trajectory, we calculated the Pearson correlation coefficient between the Euclidean distances of spatial coordinates and Euclidean distances of pseudotime values across all possible pairs of cells within the sub-trajectory. The partition score is defined as the average of the *k* Pearson correlation coefficients.

#### Differential gene score in mouse E14.5 dorsal midbrain data

We first identify genes that are differentially expressed between the RGC to NeuBs trajectory and the RGC to GlioBs trajectory. We constructed two trajectories. The first trajectory consists of all RGCs and NeuBs and the second trajectory consists of all RGCs and GlioBs. Similar to the differential test in Paella, we fit two GAM models. The null model only allows one curve shared by both trajectories (*g∼ s*(*t, k* = 3)) and the full model allows two curves for the two trajectories (*g∼ α* + *s*(*t, by* = *α, k* = 3)). *P*-values were obtained by likelihood ratio tests comparing the null and full models, and were further adjusted for multiple testing using the BH procedure^49^. Genes with FDR < 0.05 are considered as differential genes between the NeuB and GlioB trajectories.

The differential gene score is defined as the area under ROC curve where the *p*-values obtained by Paella or the random permutation approach are used to predict the differential genes between NeuB and GlioB trajectories.

#### Partition agreement in mouse olfactory bulb data

We used entropy of cluster accuracy (Hacc) and entropy of cluster purity (Hpur) to compare the spatial trajectory partition by Paella or the random permutation approach with the manually annotated partition. The details of Hacc and Hpur definitions were described in a previous study^57^. The partition agreement is defined as the sum of Hacc and Hpur. A lower partition score means a better agreement with the manual annotations.

#### Overlap score in GBM and breast cancer data

For each TCGA^32^ cancer type, we calculated the mean and standard deviation of gene expression for each gene across all samples belonging to that cancer type. Genes with zero expression across all samples were filtered out. A loess curve is then fitted where the log-transformed standard deviation of gene expression is the response variable and the mean of gene expression is the independent variable. The residual of each gene was obtained. A *t*-test was performed comparing the residuals of all genes with significant spatial differential patterns identified by Paella (or the random permutation approach) and the residuals of all non-differential genes. The overlap score is defined as the *t* statistic of the *t* test.

The same procedure was performed to obtain overlap score for each breast cancer subtype.

### Identifying subtypes of the human breast cancer spatial sample

The raw count matrix of the spatial sample were aggregated across spatial spots to create a pseudobulk gene expression vector. The pseudobulk counts were divided by the total library size of the pseudobulk, multiplied by 10^6^, and log2 transformed after adding a pseudocount of 1. The pseudobulk sample was then combined with all TCGA^32^ breast cancer samples to form one gene expression matrix, and quantile normalization^58^(`preprocessCore::normalize.quantiles()`) was performed across all TCGA^32^ samples and the spatial pseudobulk sample. Genes with larger than 0.1 expression in at least 10% of all samples were retained. For each pair of breast cancer subtypes, Limma^59^ was performed to identify top 100 differential genes with smallest FDR between the two subtypes. Principal component analysis was performed only using the union of all the top 100 differential genes across subtype pairs, and gene expression was scaled for each gene across samples. The subtype of the spatial sample was visually identified by comparing its location with the data cloud formed by different subtypes on the PCA plot.

### Testing overlap of Paella differential genes between GBM and breast cancer data

First, we calculated the number of significantly differential genes by Paella that are identified in both GBM and breast cancer data. Then we permuted the FDR output by Paella differential tests separately in GBM and breast cancer datasets and recalculated the number of shared differential genes. The permutation was performed 10, 000 times to get a null distribution of the number of shared genes. The empirical null distribution was approximated by a normal distribution where the mean and standard deviation were estimated from the empirical null distribution. The *p*-value was calculated as the tail probability of the normal distribution larger than the real number of shared genes.

## Availability of data and materials

The data used in this analysis are all publicly available. All data are described in Methods Section. The Paella package with detailed user manual is publicly available at https://github.com/Winnie09/Paella. The R package `ggplot2`^60^ for data visualization was used.

## Acknowledgments

Z.J. was supported by the National Institutes of Health under Award Number 1U54AG075936-01. W.H. was supported by the National Human Genome Research Institute of the National Institutes of Health under Award Number R00HG011468. The content is solely the responsibility of the authors and does not necessarily represent the official views of the National Institutes of Health.

## Author contributions

All authors conceived the study, developed the algorithm, conducted the analysis, and wrote the manuscript.

## Ethics approval and consent to participate

Not applicable.

## Consent for publication

Not applicable.

## Open Access

This article is distributed under the terms of the MIT License, which is a permissive license with conditions only requiring preservation of copyright and license notices.

## References

1. Qiu, X. et al. Reversed graph embedding resolves complex single-cell trajectories. Nature methods 14, 979–982 (2017).

2. Cao, J. et al. The single-cell transcriptional landscape of mammalian organogenesis. Nature 566, 496–502 (2019).

3. Ji, Z. & Ji, H. Tscan: Pseudo-time reconstruction and evaluation in single-cell rna-seq analysis. Nucleic acids research 44, e117–e117 (2016).

4. Street, K. et al. Slingshot: cell lineage and pseudotime inference for single-cell transcriptomics. BMC genomics 19, 1–16 (2018).

5. Haghverdi, L., Büttner, M., Wolf, F. A., Buettner, F. & Theis, F. J. Diffusion pseudotime robustly reconstructs lineage branching. Nature methods 13, 845–848 (2016).

6. Islam, S. et al. Quantitative single-cell rna-seq with unique molecular identifiers. Nature methods 11, 163–166 (2014).

7. Femino, A. M., Fay, F. S., Fogarty, K. & Singer, R. H. Visualization of single rna transcripts in situ. Science 280, 585–590 (1998).

8. Chen, K. H., Boettiger, A. N., Moffitt, J. R., Wang, S. & Zhuang, X. Spatially resolved, highly multiplexed rna profiling in single cells. Science 348, aaa6090 (2015).

9. Rodriques, S. G. et al. Slide-seq: A scalable technology for measuring genome-wide expression at high spatial resolution. Science 363, 1463–1467 (2019).

10. Ren, H., Walker, B. L., Cang, Z. & Nie, Q. Identifying multicellular spatiotemporal organization of cells with spaceflow. Nature communications 13, 1–14 (2022).

11. Kueckelhaus, J. et al. Inferring spatially transient gene expression pattern from spatial transcriptomic studies. BioRxiv (2020).

12. Shang, L. & Zhou, X. Spatially aware dimension reduction for spatial transcriptomics. bioRxiv (2022).

13. Pham, D. et al. stlearn: integrating spatial location, tissue morphology and gene expression to find cell types, cell-cell interactions and spatial trajectories within undissociated tissues. BioRxiv (2020).

14. Chen, A. et al. Spatiotemporal transcriptomic atlas of mouse organogenesis using dna nanoball-patterned arrays. Cell 185, 1777–1792 (2022).

15. 10x visium adult mouse olfactory bulb dataset. https://www.10xgenomics.com/resources/datasets/adult-mouse-olfactory-bulb-1-standard. Accessed: 2022-03-24.

16. Kosaka, T. & Kosaka, K. Olfactory bulb anatomy. In Reference Module in Biomedical Sciences (Elsevier, 2014). URL https://www.sciencedirect.com/science/article/pii/B978012801238304705X.

17. 10x visium human glioblastoma dataset. https://www.10xgenomics.com/resources/datasets/human-glioblastoma-whole-transcriptome-analysis-1-standard-1-2-0. Accessed: 2022-03-24.

18. Lopez-Bertoni, H. et al. Sox2 induces glioblastoma cell stemness and tumor propagation by repressing tet2 and deregulating 5hmc and 5mc dna modifications. Signal transduction and targeted therapy 7, 1–12 (2022).

19. Kupp, R. et al. Lineage-restricted olig2-rtk signaling governs the molecular subtype of glioma stem-like cells. Cell reports 16, 2838–2845 (2016).

20. Krcek, R. et al. Vascular endothelial growth factor, irradiation, and axitinib have diverse effects on motility and proliferation of glioblastoma multiforme cells. Frontiers in oncology 7, 182 (2017).

21. Chen, Z. et al. Cxcl8 promotes glioma progression by activating the jak/stat1/hif-1α/snail signaling axis. OncoTargets and therapy 12, 8125 (2019).

22. Chen, Z. et al. Prrx1 promotes stemness and angiogenesis via activating tgf-β /smad pathway and upregulating proangiogenic factors in glioma. Cell death & disease 12, 1–17 (2021).

23. Ghosh, D. et al. A cell-surface membrane protein signature for glioblastoma. Cell systems 4, 516–529 (2017).

24. Liu, J., Liu, D., Yang, Z. & Yang, Z. High lamc1 expression in glioma is associated with poor prognosis. OncoTargets and therapy 12, 4253 (2019).

25. Burton, T. et al. Bnip3 acts as transcriptional repressor of death receptor-5 expression and prevents trail-induced cell death in gliomas. Cell death & disease 4, e587–e587 (2013).

26. Treps, L., Perret, R., Edmond, S., Ricard, D. & Gavard, J. Glioblastoma stem-like cells secrete the pro-angiogenic vegf-a factor in extracellular vesicles. Journal of extracellular vesicles 6, 1359479 (2017).

27. Miyai, M. et al. Glucose transporter glut1 controls diffuse invasion phenotype with perineuronal satellitosis in diffuse glioma microenvironment. Neuro-oncology advances 3, vdaa150 (2021).

28. Pattwell, S. S. et al. A kinase-deficient ntrk2 splice variant predominates in glioma and amplifies several oncogenic signaling pathways. Nature communications 11, 1–14 (2020).

29. Hardee, M. E. & Zagzag, D. Mechanisms of glioma-associated neovascularization. The American journal of pathology 181, 1126–1141 (2012).

30. Venkatesh, H. S. et al. Electrical and synaptic integration of glioma into neural circuits. Nature 573, 539–545 (2019).

31. Takayasu, T., Kurisu, K., Esquenazi, Y. & Ballester, L. Y. Ion channels and their role in the pathophysiology of gliomas. Molecular cancer therapeutics 19, 1959–1969 (2020).

32. Weinstein, J. N. et al. The cancer genome atlas pan-cancer analysis project. Nature genetics 45, 1113–1120 (2013).

33. 10x visium human invasive ductal carcinoma breast cancer dataset. https://www.10xgenomics.com/resources/datasets/invasive-ductal-carcinoma-stained-with-fluorescent-cd-3-antibody-1-sta Accessed: 2022-03-24.

34. Zhao, E. et al. Spatial transcriptomics at subspot resolution with bayesspace. Nature Biotechnology 39, 1375–1384 (2021).

35. Debald, M. et al. Specific expression of k63-linked ubiquitination of calmodulin-like protein 5 in breast cancer of premenopausal patients. Journal of cancer research and clinical oncology 139, 2125–2132 (2013).

36. Mourskaia, A. A. et al. Abcc5 supports osteoclast formation and promotes breast cancer metastasis to bone. Breast Cancer Research 14, 1–60 (2012).

37. Hatami, R. et al. Klf6-sv1 drives breast cancer metastasis and is associated with poor survival. Science translational medicine 5, 169ra12–169ra12 (2013).

38. Zhou, F. et al. Nuclear receptor nr4a1 promotes breast cancer invasion and metastasis by activating tgf-β signalling. Nature communications 5, 1–13 (2014).

39. Sood, A. K. et al. Sam-pointed domain containing ets transcription factor in luminal breast cancer pathogenesis. Cancer Epidemiology Biomarkers & Prevention 18, 1899–1903 (2009).

40. Wu, B. et al. Mrps30-dt knockdown inhibits breast cancer progression by targeting jab1/cops5. Frontiers in oncology 9, 1170 (2019).

41. Gatti-Mays, M. E. et al. If we build it they will come: targeting the immune response to breast cancer. NPJ breast cancer 5, 1–13 (2019).

42. Boudreau, N. & Myers, C. Breast cancer-induced angiogenesis: multiple mechanisms and the role of the microenvironment. Breast cancer research 5, 1–7 (2003).

43. Li, D.-M. & Feng, Y.-M. Signaling mechanism of cell adhesion molecules in breast cancer metastasis: potential therapeutic targets. Breast cancer research and treatment 128, 7–21 (2011).

44. McInnes, L., Healy, J. & Melville, J. Umap: Uniform manifold approximation and projection for dimension reduction. arXiv preprint arXiv:1802.03426 (2018).

45. Wood, S. N., Pya, N. & Säfken, B. Smoothing parameter and model selection for general smooth models. Journal of the American Statistical Association 111, 1548–1563 (2016).

46. Dries, R. et al. Giotto: a toolbox for integrative analysis and visualization of spatial expression data. Genome biology 22, 1–31 (2021).

47. Dijkstra, E. W. et al. A note on two problems in connexion with graphs. Numerische mathematik 1, 269–271 (1959).

48. Hou, W. et al. A statistical framework for differential pseudotime analysis with multiple single-cell rna-seq samples. bioRxiv (2021).

49. Benjamini, Y. & Hochberg, Y. Controlling the false discovery rate: a practical and powerful approach to multiple testing. Journal of the Royal statistical society: series B (Methodological) 57, 289–300 (1995).

50. Zhuang, H., Wang, H. & Ji, Z. findpc: An r package to automatically select the number of principal components in single-cell analysis. Bioinformatics 38, 2949–2951 (2022).

51. Alexa, A. & Rahnenführer, J. Gene set enrichment analysis with topgo. Bioconductor Improv 27, 1–26 (2009).

52. Huang, M. et al. Saver: gene expression recovery for single-cell rna sequencing. Nature methods 15, 539–542 (2018).

53. Hafemeister, C. & Satija, R. Normalization and variance stabilization of single-cell rna-seq data using regularized negative binomial regression. Genome biology 20, 1–15 (2019).

54. Hao, Y. et al. Integrated analysis of multimodal single-cell data. Cell 184, 3573–3587 (2021).

55. Gentleman, R. C. et al. Bioconductor: open software development for computational biology and bioinformatics. Genome biology 5, 1–16 (2004).

56. Colaprico, A. et al. Tcgabiolinks: an r/bioconductor package for integrative analysis of tcga data. Nucleic acids research 44, e71–e71 (2016).

57. Hou, W., Ji, Z., Ji, H. & Hicks, S. C. A systematic evaluation of single-cell rna-sequencing imputation methods. Genome biology 21, 1–30 (2020).

58. Bolstad, B. M., Irizarry, R. A., Å strand, M. & Speed, T. P. A comparison of normalization methods for high density oligonucleotide array data based on variance and bias. Bioinformatics 19, 185–193 (2003).

59. Ritchie, M. E. et al. limma powers differential expression analyses for rna-sequencing and microarray studies. Nucleic acids research 43, e47–e47 (2015).

60. Wickham, H. ggplot2: Elegant Graphics for Data Analysis. Use R! (Springer International Publishing, Switzerland, 2016).

