## Supplementary Figure for "Decomposing spatial heterogeneity of cell trajectories with Paella"

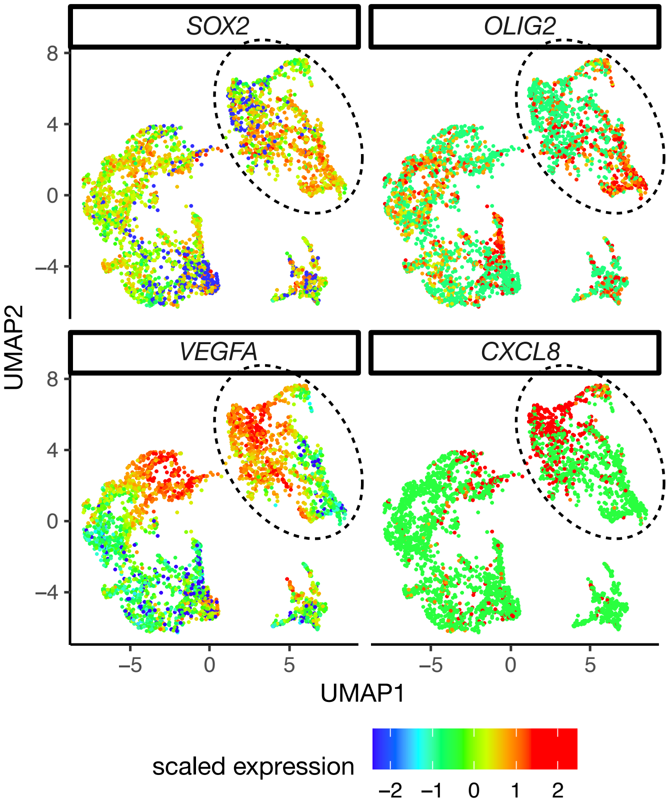


**Supplementary Figure 1.** Expression of four marker genes shown on Uniform Manifold Approximation and Projection[1] (UMAP) space in human glioblastoma dataset. Cells that used to construct pseudotime are circled in black.

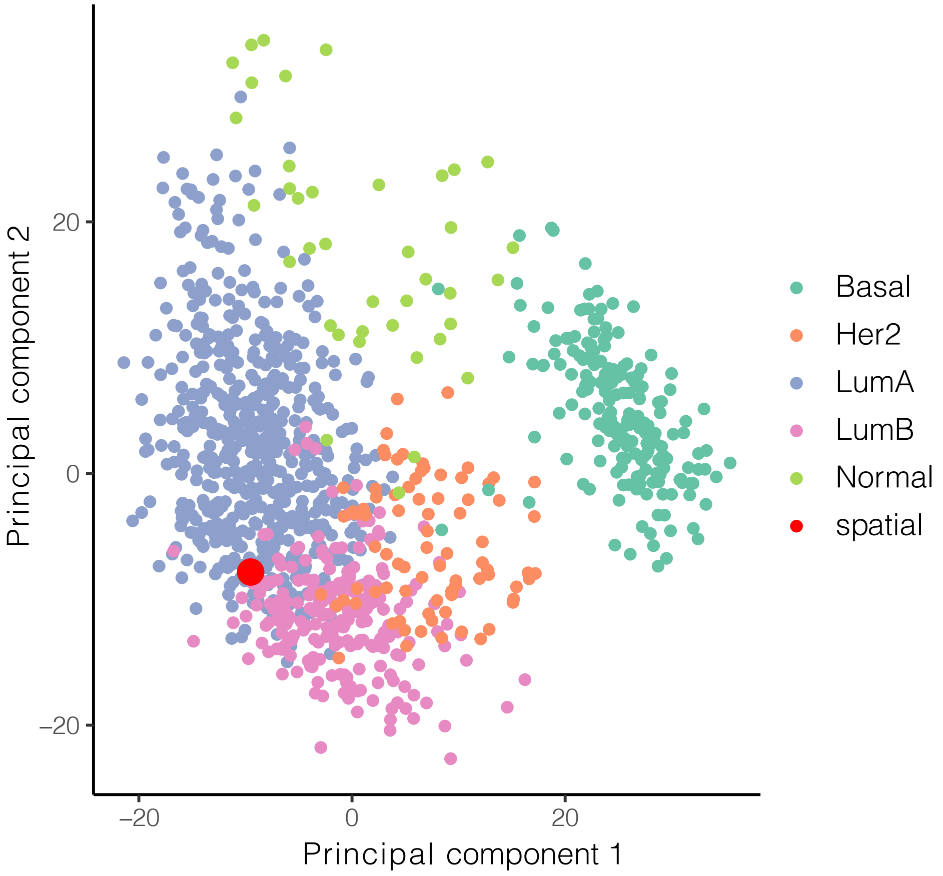


**Supplementary Figure 2.** Principal component analysis (PCA) combining the human breast cancer spatial samples and TCGA breast cancer samples of different subtypes. The spatial sample is classified as Luminal A subtype since it is within the data cloud of TCGA samples of Luminal A subtype.

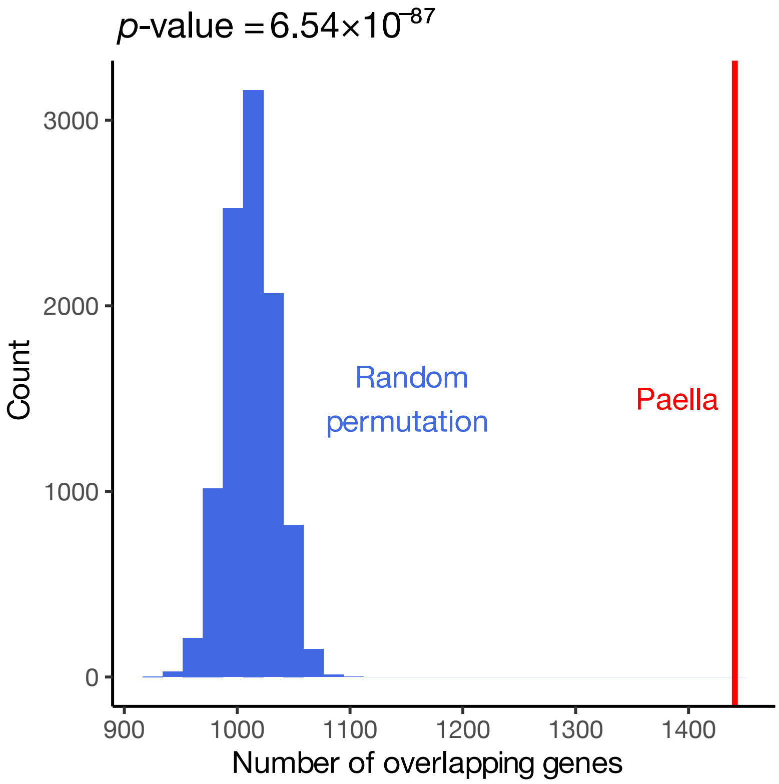


**Supplementary Figure 3.** Number of overlapping genes comparing human glioblastoma and breast cancer datasets. The number obtained by Paella is shown as a red vertical line. The numbers obtained by random permutations are shown as a blue histogram.

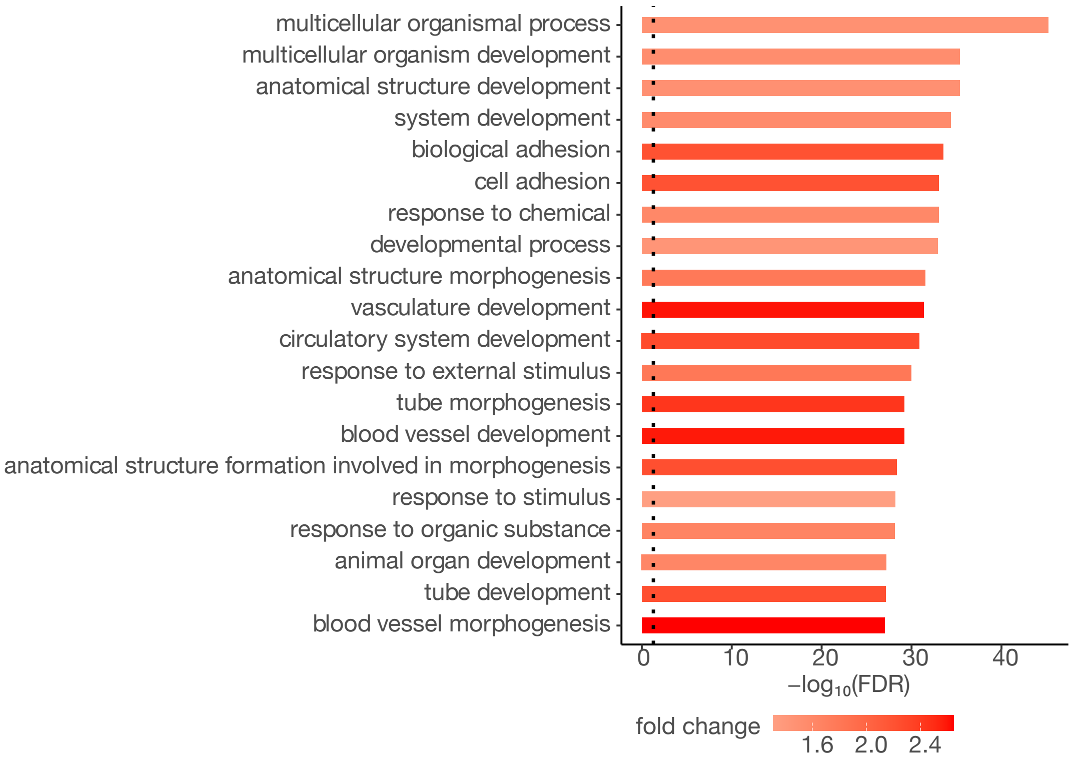


**Supplementary Figure 4.** Top enriched gene ontology (GO) terms of the genes with differential temporal patterns across spatial sub-trajectories identified by Paella shared by human glioblastoma and breast cancer datasets. An FDR cutoff of 0.05 is indicated as a vertical black dashed line.

### References

**1.** McInnes, L., Healy, J. & Melville, J. Umap: Uniform manifold approximation and projection for dimension reduction. *arXiv preprint arXiv:1802.03426* (2018).
